# In-situ spectroscopic detection of large-scale reorientations of transmembrane α-helices during viroporin channel opening

**DOI:** 10.1101/2023.06.22.546036

**Authors:** Ronja Paschke, Swantje Mohr, Sascha Lange, Adam Lange, Jacek Kozuch

## Abstract

Viroporins are small ion channels in membranes of enveloped viruses that play key roles during viral life cycles. To use viroporins as drug targets against viral infection requires in-depth mechanistic understanding and, with that, methods that enable investigations under in-situ conditions. Here, we apply surface-enhanced infrared absorption (SEIRA) spectroscopy to Influenza A M2 reconstituted within a solid-supported membrane, to shed light on the mechanics of its viroporin function. M2 is a paradigm of pH-activated proton channels and controls the proton flux into the viral interior during viral infection. We use SEIRA to track the large-scale reorientation of M2’s transmembrane α-helices in-situ during pH-activated channel opening. We quantify this event as a helical tilt from 26° to 40° by correlating the experimental results with solid-state NMR-informed computational spectroscopy. This mechanical motion is impeded upon addition of the inhibitor rimantadine, giving a direct spectroscopic marker to test antiviral activity. The presented approach provides a spectroscopic tool to quantify large-scale structural changes and to track the function and inhibition of the growing number of viroporins from pathogenic viruses in future studies.

---

Viroporins are a family of small, ion channel-forming viral membrane proteins that play essential roles at various steps of the viral life cycle and, thus, during infection.^[1]^ To date, many viroporins, such as the HIV-1 Vpu, Hepatitis p7, and SARS CoV-2 E protein, have been characterized regarding their structure, function, and virological significance,^[2]^ and new members are added steadily to this class of proteins.^[3]^ However, it was the M2 proton channel from Influenza A virus (IAV)^[4]^ that garnered much interest for this family of proteins, when it was shown that its viroporin function, and, thereby, viral infection, can be inhibited by adamantane drug binding.^[5]^ With the prevalence of diseases caused by pathogenic viruses but scarcity of potent treatments, this demonstrated that viroporins present a promising target for antiviral therapy. As such, a detailed understanding of their mechanisms of action and inhibition is required and, consequently, methods that enable their investigation under in-situ conditions.

**Scheme 1.**
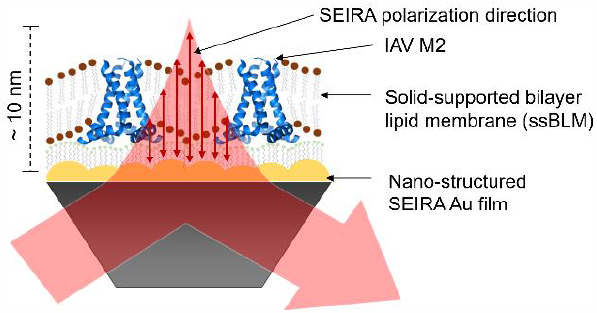
Schematic representation of the Influenza A virus (IAV) M2-containing solid-supported bilayer lipid membrane (ssBLM) on a nano-structured SEIRA Au film. SEIRA probes IR spectral changes within ca. 10 nm, i.e., of a single ssBLM layer, and provides information on reorientation events due to the polarization direction (surface-selection rule).

The IAV M2 protein is a tetrameric pH-dependent proton channel inside the viral envelope, which is activated at pH values < 7 after endocytic entry of the virion, a step required for release of genetic material into the host cell.^[6]^ Extensive studies have revealed that four pore-lining His37 and Trp41 residues near the center of the transmembrane (TM) helices are crucial in the gating and proton conductance mechanism, respectively.^[7]^ According to the accepted model, protonation of the His37 tetrad from the viral exterior side opens the channel via electrostatic repulsion between the imidazolium groups to enable proton flow into the virion.^[8–10]^ The chemical step of His37 protonation has been investigated in detail using various methods, including x-ray crystallography,^[11,12]^ solid-state nuclear magnetic resonance (ssNMR),^[13–17]^ and molecular dynamics simulations.^[18,19]^

However, quantifying the large-scale reorganization of the tetrameric structure associated with the widening of the pore radius has been much more challenging, in particular from an experimental perspective. Whereas structures of M2 show different relative alignments of the TM helices, their comparison is complicated as they refer to constructs of various lengths. As such, optical spectroscopic investigations that enable monitoring channel function in-situ have become important, as exemplified by 2D infrared spectroscopic work that proposed twisting motions of the TM helices after pH activation.^[20]^

To shed more light on the mechanics of pH-induced channel opening of M2, we performed surface-enhanced infrared absorption (SEIRA) spectroscopic measurements on M2 reconstituted within a solid-supported bilayer lipid membrane (ssBLM; Scheme 1), as a minimal model system for the viral membrane.^[21,22]^ SEIRA spectroscopy allows monitoring protonation events^[23,24]^ as well as quantifying large-scale helical reorientation^[25,26]^ in-situ upon pH activation. This is achieved using a nano-structured Au film that provides plasmonic infrared (IR) signal enhancement of the reconstituted proteins in a single ssBLM by a factor of about 10^2^.^[22,27]^ In turn, the bulk solution is spectrally invisible due to a sharp distance-dependent attenuation of the enhancement after just 10 nm from the surface. Furthermore, the surface selection rule enables detecting molecular (re)orientation events, as only changes in IR modes with dipole moments perpendicular to the surface are enhanced. Here, we quantify the SEIRA-based results by comparing them to density functional theory (DFT)-derived computational spectroscopic predictions that are informed by structural data from ssNMR spectroscopy.

The IAV M2_18-60_ construct used in this work is composed of a short segment of the unstructured N-terminal domain (res. 18 -21), the TM helix (res. 22 -46) and a segment of the short amphipathic helix (res. 47 -60),^[28]^ which was expressed in *E. coli* and reconstituted into 1,2-diphytanoyl-sn-glycero-3-phosphatidylcholine (DPhPC) liposomes according to Fu et al.^[29]^ The M2-proteoliposomes were deposited on a 6-mercaptohexanol-functionalized SEIRA Au surface to create a M2-ssBLM system revealing characteristic bands at 1659 cm^-1^ and 1547 cm^-1^ that show an isotopic downshift by -45 to -25 cm^-1^ upon global ^13^C/^15^N-labelling of the M2 protein. These signals can, therefore, be assigned to the amide I and amide II modes of the α-helical protein backbone structure (Figure 1A, top).^[30]^ The peaks at 1736 and below 1500 cm^-1^ on the other hand do not shift and can, thus, be ascribed to the lipids of the ssBLM.^[25,31]^

**Figure 1.**
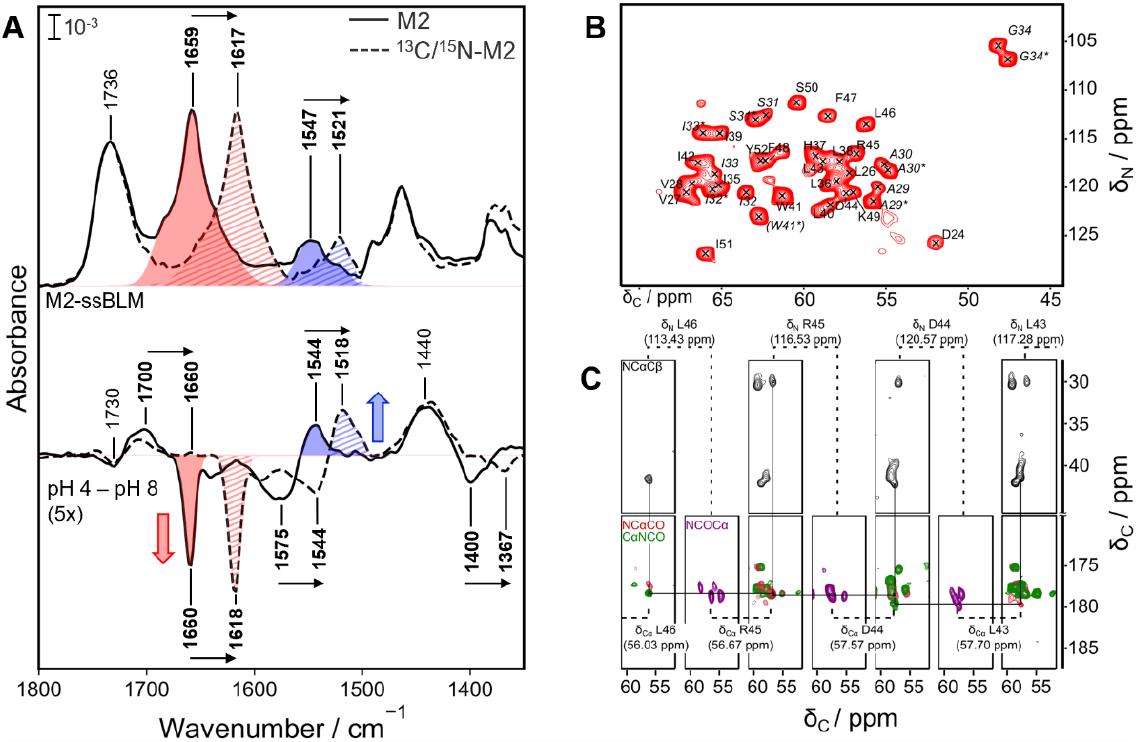
A: SEIRA spectra of IAV M2 within ssBLM (top) and difference spectra after changing the pH from 8 to 4 (bottom) for unlabeled M2 (solid lines) and _13_C/_15_N-labelled M2 (dashed lines); isotope shifts are shown by arrows. Amide I and II bands are indicated in red and blue, respectively; red and blue arrows highlight the direction of the corresponding difference peaks. B: 2D N-Cα correlation spectrum of _13_C/_15_N-M2 in DPhPC proteoliposomes at pH 7.8. Assigned resonances are indicated in the spectra. In the N-Cα spectrum, carbon and nitrogen resonances of the assigned peaks correspond to the same residue. Doubled peaks are indicated in italics, with the second resonance marked with an asterisk (*). W41* could not be assigned unambiguously from our data, thus being in brackets. C: Sequential walk for the amino acid stretch L46 to L43 shown in a strip plot of 3D experiments. Spectra are shown in black (NCαCβ), red (NCαCO), green (CαNCO) and purple (NCOCα), strips sharing the same Cα- or N-chemical shift are marked with dashed lines and the respective residue number, steps of the backbone walk are indicated with solid lines.

To assure that these amide I/II signals reflect a well-folded protein, we recorded ssNMR spectra on the same proteoliposomes and demonstrate the native fold of M2 with most of the residues in the TM domain and the amphipathic helix being well resolved in a ^15^N-^13^C fingerprint spectrum (Figure 1B). Assignments (see Figure 1C for a sequential backbone walk) are in line with previous studies^[32]^ and suggest a tetrameric form, with peak doubling occurring for res. 28 to 34, pointing to structural plasticity in this region. Residues at the N-terminal part (res. 18 -23) as well as at the C-terminal end (res. 53 -60) could not be detected in dipolar coupling-based experiments, probably due to higher flexibility in these domains.

To gain information on the pH-dependent mechanism of the channel, SEIRA difference spectra were recorded after decreasing the bulk solution’s pH from 8 to 4 (Figure 1A, bottom). Difference spectroscopy is a common tool in IR spectroscopy as it allows to focus on changes upon a stimulus, while ignoring the portion of the signal that remained unchanged. As such, negative and positive peaks (denoted as “-“ or “+“, respectively, in the following) are ascribed to species that have been lost or gained during the pH change, respectively. Accordingly, we note peaks at 1730 and at 1460 cm^-1^, which are not affected by the ^13^C^15^N-isotope labelling of M2 and, therefore, can be assigned to C=O stretching of the lipid’s ester group^[33]^ and CH_x_ deformations of the lipid’s acyl chains, respectively.^[34]^ However, we also recognize prominent signals that shift upon ^13^C/^15^N-labelling and are thus caused by changes of M2. First, signals at 1700(+), 1575(-), and 1400(-) cm^-1^ were detected (~1660, 1544, and 1367 cm^-1^, respectively, in ^13^C/^15^N-M2), which are clear markers for the protonation of carboxylate groups.^[35]^ The M2_18-60_ construct contains multiple carboxylate groups, namely at Asp21, Asp24 or Asp44; a potential protonation of one of these residues was suggested previously to contribute to the channel conductivity.^[36,37]^ Second, a prominent couple of a negative peak at 1660(-) and a positive signal at 1544(+) cm^-1^ is observed (1618 and 1517 cm^-1^ in ^13^C/^15^N-M2), which can be assigned to the amide I and II vibrations of the α-helical backbone of M2. Intriguingly, this specific feature of amide I and II peaks with intensity of opposite sign is a clear IR spectroscopic marker of reorienting α-helices, as observed in our previous work on antimicrobial peptides.^[25]^ This spectroscopic feature relies on the approximately perpendicular orientation of the transition dipole moments (TDMs) of the amide I and II vibrations (mostly C=O stretch and N-H bend of the peptide groups, respectively – see below and scheme in Figure 2B). In combination with the surface selection rule in SEIRA, the orientation of the TDMs results in a preferential enhancement of one of the two vibrational modes. In the present case, this effect provides us with direct insight into the reorientation of the α-helices of M2 during pH-activated channel opening. We would like to note that we do not detect the protonation of the pH-sensing His37 residues, since the imidazole sidechain generally does not provide a clear IR-spectroscopic marker.^[38]^

**Figure 2.**
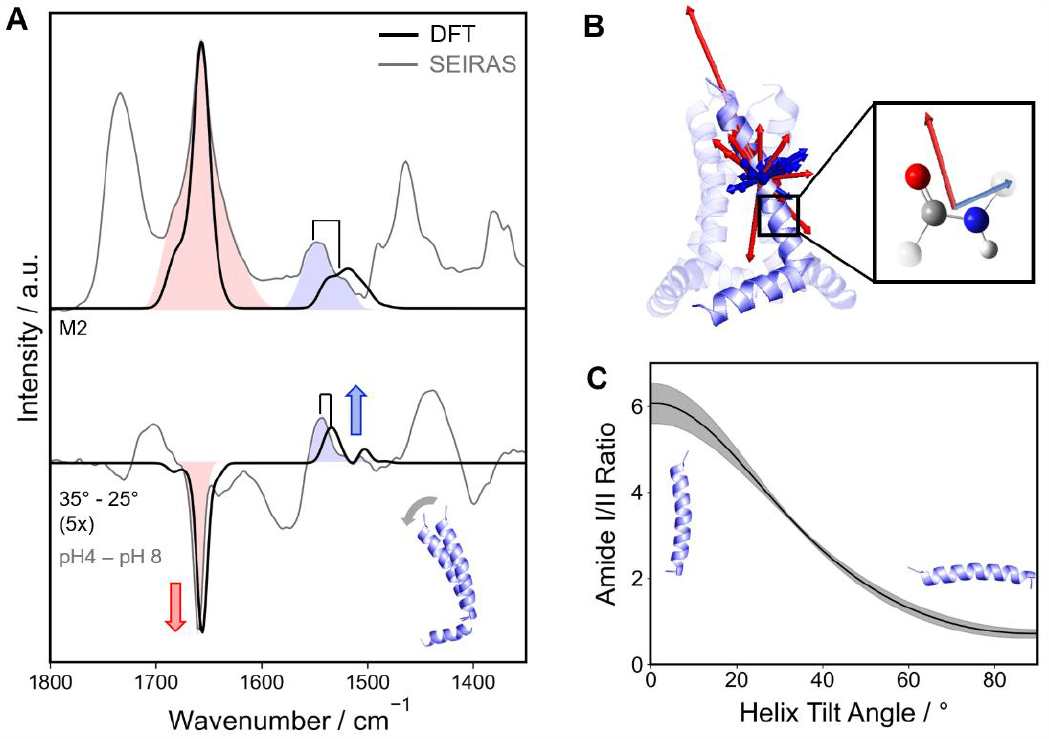
A: DFT-computed spectra (FWHM set to 12 cm^-1^) of the M2 protein backbone (black, top) and difference spectra for a change in helix tilt angle from 25° to 35° (black, bottom) – see inset. For comparison, light colors indicate the related experimental spectra; amide I and II bands are highlighted in red and blue, respectively. Spectra were normalized to the amide I peak for clarity. B: The TDM vectors of amide I (red) and amide II (blue) of a TM α-helix in M2 are aligned preferentially along or perpendicular to the helix axis (PDB: 2l0j), respectively. The preferential orientation originates from coupling between the single amide units, whose amide I and II TDM vectors are approximately perpendicular to each other. C: Trend of the amide I/II ratio of the peak areas dependent on the helix tilt angle from 0° to 90° in respect to the membrane surface normal. Confidence is obtained from different starting structures in the DFT calculation (Figure S2).

To illustrate this orientation-dependent spectroscopic effect more clearly and demonstrate its usefulness for a quantitative analysis of the orientation of α-helices, we computed the theoretical spectra of the protein backbone with density functional theory (DFT) using the approach previously reported in Forbrig et al.^[25]^ and related work by Kubelka et al.^[39]^ Accordingly, in a simple model (Figure 2B), we performed a geometry optimization and normal mode analysis of the TM helix (res. 22 -46) and the amphipathic helix (res. 48-60) individually. We used several starting structures based on the ssNMR-determined dihedrals obtained in this work (Table S1), a previously reported ssNMR structure (pdb 2l0j),^[40]^ and a fully unconstrained model giving overall similar results as summarized in Figure S2. Based on the TDMs of the normal modes contributing to the amide I and II bands (Figure 2B), we can model the polarized spectra. Figure 2A (top) shows a representative resulting computational absolute spectrum of the amide I and II signals (set to FWHM of 12 cm^-1^) by considering the average orientation of the TM helices of 25° with respect to the symmetry axis of M2 as reported in literature for the closed state (SI Table 2). As in our previous work, the computed amide I/II-spectrum matches the experimental one very well in intensity and position. Only the amide II shows a slight shift by approximately 10 cm^-1^, which is acceptable considering the level of theory used in this work (bp86/6-31g*).^[25,39,41,42]^ Modelling a rigid body reorientation of the TM helix, we determined the computational difference spectra considering the polarization direction according to the surface selection rule in SEIRA (Figure 2A, bottom; experimental data are shown in light colors for comparison). We see a striking similarity in the spectral features of the amide I peak at 1659(-) cm^-1^ and the amide II region at 1559(+) to 1492(+) cm^-1^ between theoretical and experimental SEIRA difference spectra, providing the direct proof-of-concept for the experimental detection of α-helical reorientation upon pH-activated M2 channel opening. It is important to mention that a reorientation of the amphipathic helix (SI Figure S1D) shows a distinctly different spectral pattern, such that this approach suggests that the reorientation is localized to the TM region.

Having demonstrated the α-helical reorientation qualitatively, we now have a tool to monitor the orientation of the TM helices of M2 via changes in the amide I and II intensities. More specifically, we can use the amide I/II ratio as a measure for the quantification of the absolute orientation. We calculated the DFT-based amide I/II intensity ratio based on absolute spectra (SI Figure S1 A, B) for TM helix tilt angles from 0° to 90° with respect to the surface normal (Figure 2C). To estimate the accuracy of the “amide I/II ratio vs. angle” calibration, we show the average and standard deviation over all structural models shown in Figure SI2. We see that for an ideal vertical orientation of the TM helix in the M2 channel, a ratio of ~6 is expected, while a horizontal orientation would provide a ratio of ~0.7. Previously, we have shown that ideal vertical orientation of a helix can provide amide I/II ratios of > 15.^[21,25]^ In the present case, the amphipathic helix is oriented horizontally, where a amide I/II ratio of around 1 is expected,^[43]^ lowering this maximum achievable value.

Turning back to the experimental results, we now aim to quantify the pH-dependent reorientation of M2’s TM helices during channel opening at pH < 7. To monitor the opening of the M2 channels within the ssBLM, we performed a pH titration from 8 to 4 (Figure 3A, top), which shows a pH-dependent increase of the spectral features observed above. As shown in Figure 3A (middle), this trend matches very well the DFT-based spectra following a reorientation in a range of 26 to 40°. Inspecting the changes of amide I and II, we notice that the absorption intensities respond to two pK_A_ values, one at 7 and one at or below 5.5 based on a fit of two coupled sigmoidal curves (Figure 3B, SI Figure S1C includes fits of the markers for the carboxylate protonation). Previous work has suggested that in the closed state of M2 at least one His residue of the His37-tetrad is protonated, and further protonation occurs with pK_A_ values of 7.1 and 5.4, respectively.^[17]^ Despite not detecting the direct IR fingerprint of the imidazole residues, we can observe the same protonation transitions as part of the amide I and II changes, and are, therefore, consistent with previous work. Our data further suggests that the more acidic pK_A_ does not originate (solely) from the protonation of the His37-tetrad but is coupled to aspartic acids (SI Figure S1 C). The pH-dependent amide I/II ratio (Figure 3C) obtained from the integrated absolute peak areas, changes from about 4.1 to 2.5 in the explored pH range. Using the calibration from computational spectra (Figure 2C), we are now able to translate this data to a pH-dependent TM helix tilt angle (Figure 3D). We obtain an angle of 26° at pH 8, which upon acidification results in a helix tilt of up to 40° and, therefore, an increase in angle by 14°. This result is consistent with literature where the average TM helix angles of the solved structures is 25° (see Table S2) and a 10 -12° difference for an opened and a closed state of the proton channel has been suggested previously.^[11,12]^ Interestingly, our data further suggests that this mechanical opening is guided by the more acidic pK_A_ value, as a plateau region is observed in the pH region of 8 to 7. It is important to note that the previously suggested pH-induced twist of the helices is not contradicted by our results and may be part of the mechanics of channel opening.^[20]^

**Figure 3.**
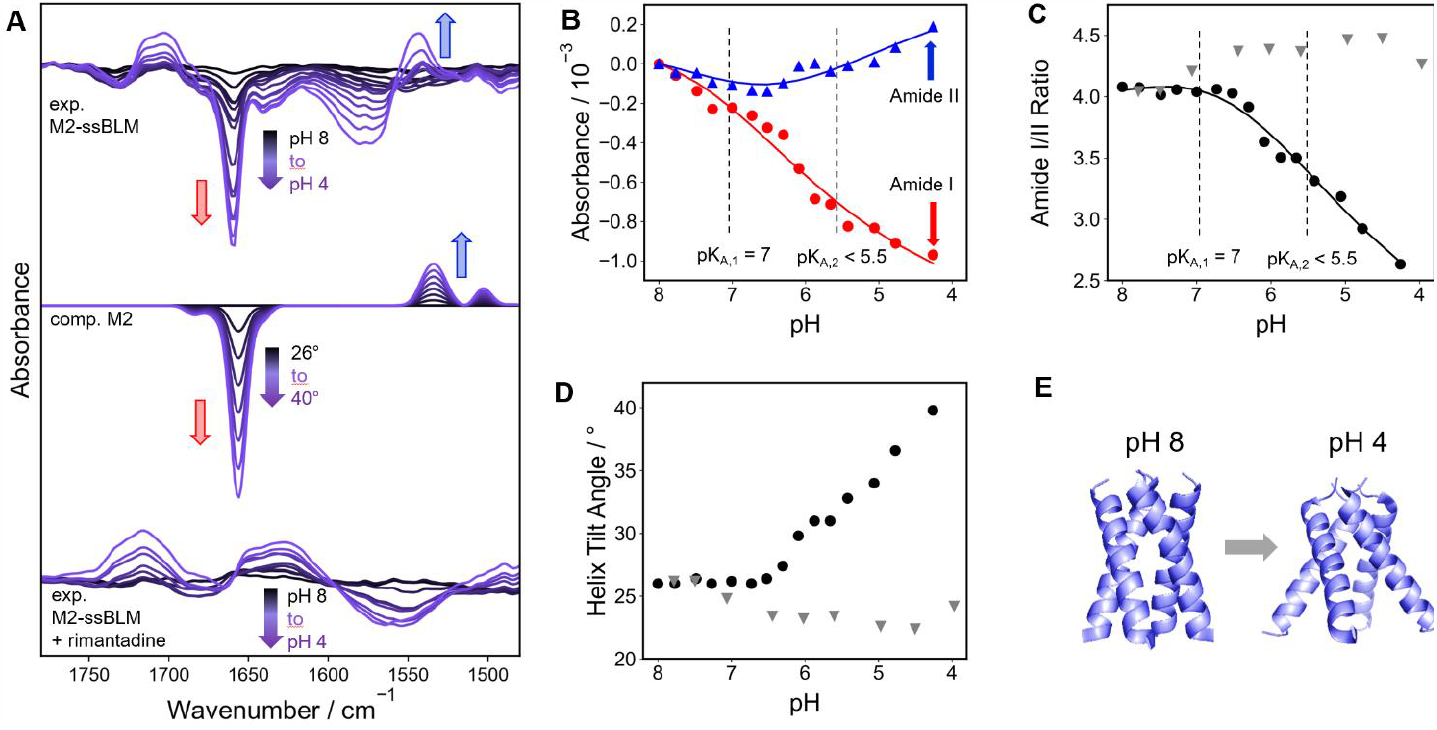
A: Experimental difference spectra of the pH titration of M2 in a ssBLM (top) and computational difference spectra for a stepwise TM helix reorientation from 26° to 40° (middle); experimental pH titration after adding the inhibitor rimantadine (bottom). B: Experimental changes in the pH-dependent absorbance of amide I at 1559 cm^-1^ (red) and the amide II at 1544 cm^-1^(blue). The fitted lines and pK_A_ values were obtained from global double sigmoidal fits considering pH-dependent spectral changes with two pK_A_ values; pK_A2_ is indicated as lower as 5.5 since the plateau at acidic conditions was not fully resolved. C: Experimental amide I/II ratio without inhibitor (black) and after adding rimantadine (grey triangles) over pH. The fitted lines result from intensity changes in B. D: Average TM α-helix tilt angles of M2 without inhibitor (black dots) and after adding rimantadine (grey triangles) depending on pH, calculated considering the computational calibration curve in Figure 2C. E: Schematic representation of the structure of M2 before (PDB: 6bkl) and after (PDB: 6boc) pH-dependent channel opening according to SEIRA spectra.

Finally, having determined the IR spectral features of channel opening, we repeated the SEIRA pH titration in the presence of the known M2 inhibitor rimantadine (Figure 3A, bottom). Interestingly, while still observing the protonation of aspartates, the opposing changes in the amide I and II bands are notably absent. Plotting the amide I/II ratio and the associated helical tilt angles over pH (Figure 3C, D), we can determine similar angles in the closed state of around 26°, but the large changes of the tilt angle of the TM helix at lower pH are absent. This provides direct evidence for a model, in which (part) of the inhibition mechanism involves a blockage of the dynamical changes of helix alignment.

To conclude, with the presented approach we have a tool to track in-situ the large-scale structural changes associated with M2’s pH-dependent channel activation, an essential process in IAV infection. Combining the surface-selection rule inherent to SEIRA spectroscopy and ssNMR-informed computational spectroscopy, we quantify these changes as a reorientation of the channel’s transmembrane helices by 14° (Figure 3E). However, this mechanical opening is inhibited upon addition of the antiviral drug rimantadine. As such, our data confirms the plethora of structural data that proposed that “fanned out” structures resolved an opened conformation of the channel, which we observed here under operating conditions. In the future, we aim to utilize this approach to enable a combined structural and functional analysis of viroporins of current relevance and contribute to the discovery of new inhibitors. In particular, combining results from SEIRA spectroscopy and ssNMR, which visualize complementary structural changes, can be of high benefit towards an extensive understanding of the action of viral proteins and finding of novel antiviral drugging strategies.

## Supporting Information

Materials and methods (sample preparation, SEIRA spectroscopy, ssNMR spectroscopy, DFT calculations) and supplementary figures and tables. The authors have cited additional references within the Supporting Information.^[44–52]^

## Author Contributions

R.P. and S.M. contributed equally to this work.

## Acknowledgements

R.P. and J.K. thank Kenichi Ataka (Freie Universität Berlin, Germany) for helpful discussions and Joachim Heberle (Freie Universität Berlin, Germany) for the continuous support. We thank James Chou (Harvard Medical School, Boston, MA, USA) for generously providing the expression plasmid coding for the IAV M218-60 construct. This work was supported by DFG via the Sonderforschungsbereich 1078 “Protonation Dynamics in Protein Function” (project number 221545957, projects B09 and B10; J.K. and A.L., respectively).

