## Supplementary information for "In-situ spectroscopic detection of large-scale reorientations of transmembrane α-helices during viroporin channel opening"

### Materials and Methods

#### Sample Preparation

The IAV $M2_{18\text{-}60}$ construct was expressed into inclusion bodies as C-terminal fusion with the $(His)_9$-trpLE polypeptide by overexpression in E. Coli Bl21(DE3) as previously described[1] with slight modifications. In brief: A single colony of freshly transformed E. Coli Bl21(DE3) was used to inoculate 5 ml of LB medium for 2-3 hours at 37°C. 2 ml of this preculture was used to inoculate 200 ml of LB medium. After incubation overnight at 30°C, 2 l of fresh LB medium was inoculated with this culture to an OD of 0.2. After incubation at 37°C until an OD of 0.6 was reached, the temperature was lowered to 22°C and expression was induced 30 min later by adding 0.5 mM IPTG. For the expression of $^{13}C/^{15}N$-labelled IAV $M2_{18\text{-}60}$ D-Glucose-$^{13}C_6$ was used as the sole carbon source and $^{15}NH_4I$ as the sole nitrogen source. Bacteria were harvested after 18 h by centrifugation and stored at -80°C.
Frozen cell pellets were resuspended in 200 ml $H_2O$ and disrupted by sonication (40% power, 3 intervals of 1 min). Inclusion bodies were isolated by centrifugation at 16,000 rcf for 20 min and washed twice in 40 ml $H_2O$ containing 2% Triton-X100. After washing the inclusion bodies again in $H_2O$, the remaining pellet was resuspended in 200 ml of 6M guanidine hydrochloride (GuHCl) containing 10 mM imidazole and applied to a 5 ml HisTrap™ HP (Cytiva) column at a flow rate of 3 ml/min. The column was washed with 5 column volumes of 6M GuHCl with 10 mM imidazole and the protein was eluted in column volumes of 6M GuHCl with 1M imidazole. The flow-through of the loaded sample was combined with the wash fraction and reloaded onto the column, followed by another elution with 5 column volumes. Both elution fractions were pooled, and the buffer was changed to 0.2 M HCl in 6M GuHCl.

Cleavage of the trpLE polypeptide was carried out in the dark and under a light stream of nitrogen after addition of 4 to 5 g cyanobromide (CNBr). The reaction was stopped after 3 h by dialysis against $H_2O$, and the cleaved protein was lyophilized and dissolved in 6M HCl. The M2 pore domain was further purified by reversed-phase chromatography using a Zorbax SB-C3 column on a AZURA-Bio HPLC (KNAURGermany). HPLC fractions were analyzed by MALDI-TOF and the fractions containing M2 were pooled and lyophilized.

Lyophilized M2 and DPhPC lipid (1:1 w:w ratio, i.e., 1:6 mol:mol ratio) was added to denaturing buffer (6M guanidine, 40 mM phosphate, 30 mM glutamate, 3 mM sodium azide, pH 7. 8, >=33mg/mL OG detergent) and dialyzed against 1 L sample buffer (40 mM phosphate, 30 mM glutamate, 3 mM sodium azide, pH 7.8) for 7 days with 2 dialysis buffer changes per day.

#### SEIRA spectroscopy

SEIRA spectroscopy was performed with a Kretschmann-ATR configuration in a Bruker Vertex80v spectrometer. A nano-structured SEIRA gold film was deposited on a silicon ATR-IR element by electroless deposition.[2] All spectra were recorded in a spectral range of 4000 to 1000 $cm^{-1}$ by accumulating 200 Scans with a resolution of 4 $cm^{-1}$. A liquid $N_2$-cooled photoconductive MCT detector was used.

The solid-supported lipid bilayer was constructed by first adding 0.1 M 6MH in isopropanol on the nano-structured gold surface to form a self-assembled monolayer overnight. After rinsing with isopropanol and buffer, the M2-DPhPC proteoliposomes were added. The deposition process was monitored using SEIRA spectroscopy. After equilibration and rinsing with buffer and water, pH difference spectra were first recorded by exchanging 100 mM Phosphate buffer of pH 8 and of pH 4 repeatedly. 100 mM phosphate buffer pH 8 or 150 mM NaCl, 20 mM MES, 20 mM HEPES buffer was then recorded as the reference for the pH titration experiment (both buffering conditions gave similar results). Adding 1 M HCl to achieve 0.2 to 0.5 increments of pH steps was monitored by the pH meter and by recording SEIRA spectra until equilibrated for every pH step.

#### Solid-state NMR spectroscopy

For solid-state NMR experiments, the fully $^{13}C^{15}N$-labeled sample was packed into a 3.2 mm rotor. Solid-state NMR measurements were performed on a 700 MHz Bruker NMR spectrometer equipped with a 3.2 mm triple-resonance Efree magic-angle spinning (MAS) probe (Bruker BioSpin). All spectra were recorded at 17.5 kHz spinning rate and approximately 15 °C as determined from the water signal chemical shift. DSS (4,4-dimethyl-4-silapentane-1-sulfonic sodium salt) was used as an external chemical shift reference.[3] 90° pulse radio frequency amplitudes were set to 83.3 kHz for protons, 50 kHz for carbons and 35.7 kHz for nitrogens. During evolution and detection periods, high power proton decoupling using the SPINAL-64 pulse sequence was employed.[4] For the shown cross polarization (CP)-based 2D $^{15}N$-$^{13}C\alpha$ correlation spectrum, pulse powers were optimized around 5 kHz (nitrogen) and 13 kHz (carbon) and the CP contact time was set to 4.3 ms.

The chemical shift assignments were performed as described in previous works.[5] In short, a set of six 3D-experiments, NCOCα, NCαCO, CαNCO, NCαCB, NcoCαCB and NCOcαCB, was recorded, enabling the assignment of backbone atoms from most of the M2 residues. All spectra were processed in TopSpin 4.1.1[6] and analyzed using CCPNMR3.1.0[7] software. Protein backbone torsion angles used in the DFT calculation were predicted from the assigned chemical shifts shown in table S3 using the TALOS+ software in no-Proton mode.[8]

#### Density Functional Theory Calculations

The computation of the theoretical SEIRA spectra were carried out in accordance to the work by Forbrig et al. (2018)[9]. Gaussian 16[10] was used to perform geometry optimization and normal mode analysis on the BP86[11,12] level of theory with the 6-31g* basis set and the polarizable continuum model simulating water as a solvent.[9,13] The ssNMR structure of the M2 (22-62) construct solved by Sharma et al. (2010)[14] (PDB ID: 2l0j) was used as a starting structure. The construct consists of two alpha helical parts that were considered separately. The amino acid sequence from 22 to 47 was considered as the transmembrane helix, while amino acids 49 to 61 were considered as the 2$^{nd}$ helix. The protein backbone was extracted from the PDB file and the dihedral angles were either left without restrains or they were restrained in analogy to Keiderling et al[15]. Restraints were applied for both conformations that were solved from the assignments of the ssNMR spectra (Table S3, Table S1) and for comparison they were also restrained with the dihedral angles solved by Sharma et al (2010)[14]. The amino acid residues were removed to leave the protein as an alanine peptide, thus further facilitating the calculation. The transition dipole moments were extracted from normal mode analysis and used to reproduce the theoretical spectra for different orientations of the helices. Calculation of the spectra considering the TDMs and their projection onto the axis perpendicular to the SEIRA surface and simulation of the reorientation were carried out using home-built python scripts. Difference spectra were calculated with the initial orientation of both helices as extracted from the pdb-file as the reference absolute spectrum. The opening trajectory was extracted from the x-ray structures solved by Thomaston and Degrado (2019) (PDB ID: 6mjh).[16]

### Supplementary Figures and Tables

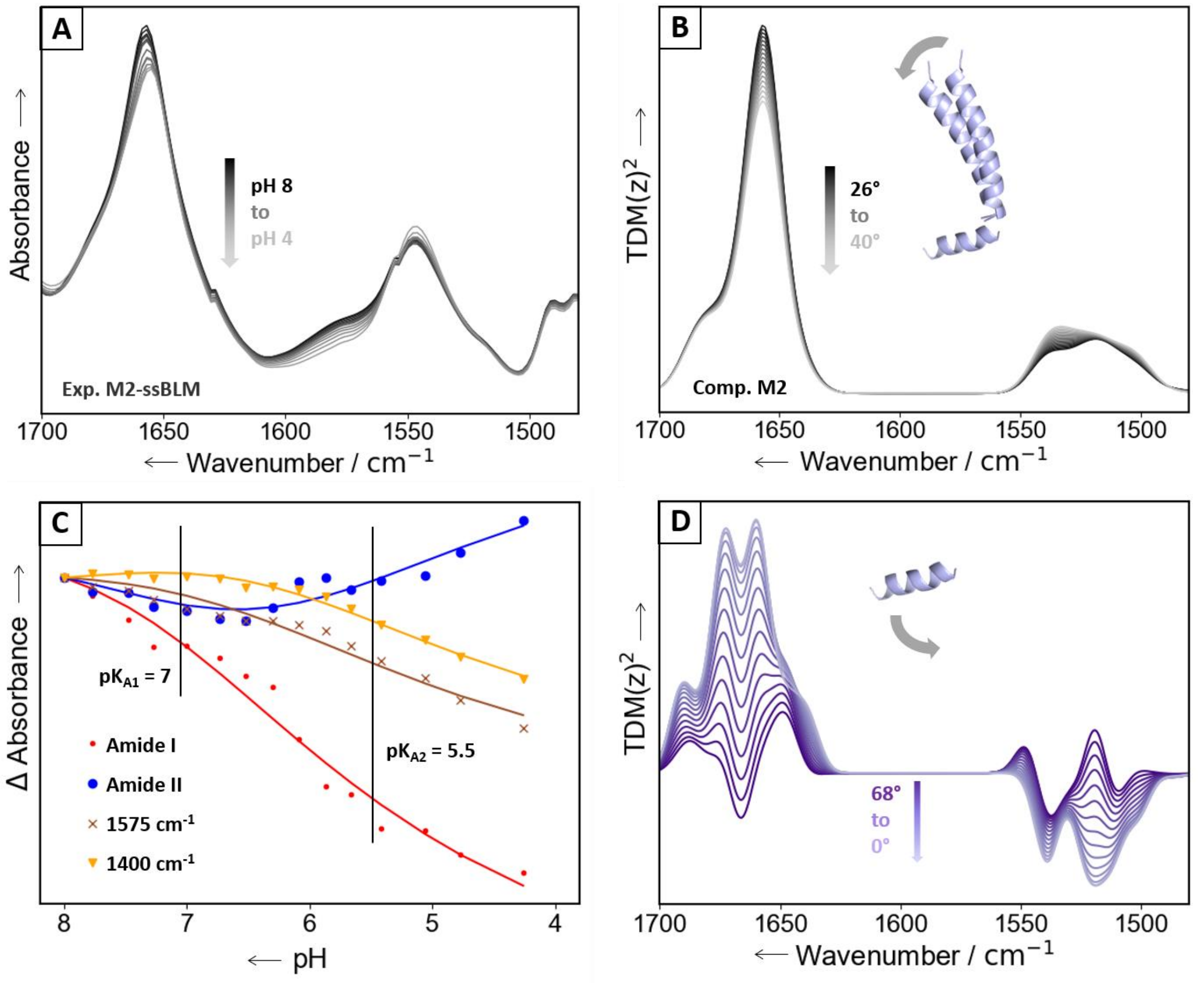

Fig S1: A: *SEIRA absolute spectra of the sample at different pH between pH 8 and 4. Areas of the amide bands were calculated to obtain the amide I/II ratios that were then used for the helix tilt angle quantification. B: Computational absolute spectra of the protein (composed of both helices) for a steady change in helix tilt angle from 26 ° to 40 ° of the TM helix. Areas of the amide bands were calculated to obtain the amide I/II ratios. C: pH-dependent changes in amide I, II and carboxylate peak intensities. A global, coupled double sigmoidal fit reveals two pK values (at 7 and at < 5.5). D: Computational difference spectra obtained from the TDMs of the amphipathic helix when simulating a reorientation from the original orientation of 68° to 0° from the surface normal. The spectra show no similarity to experimental results, suggesting that the opening mechanism does not involve a large reorientation event of the amphipathic helix.*

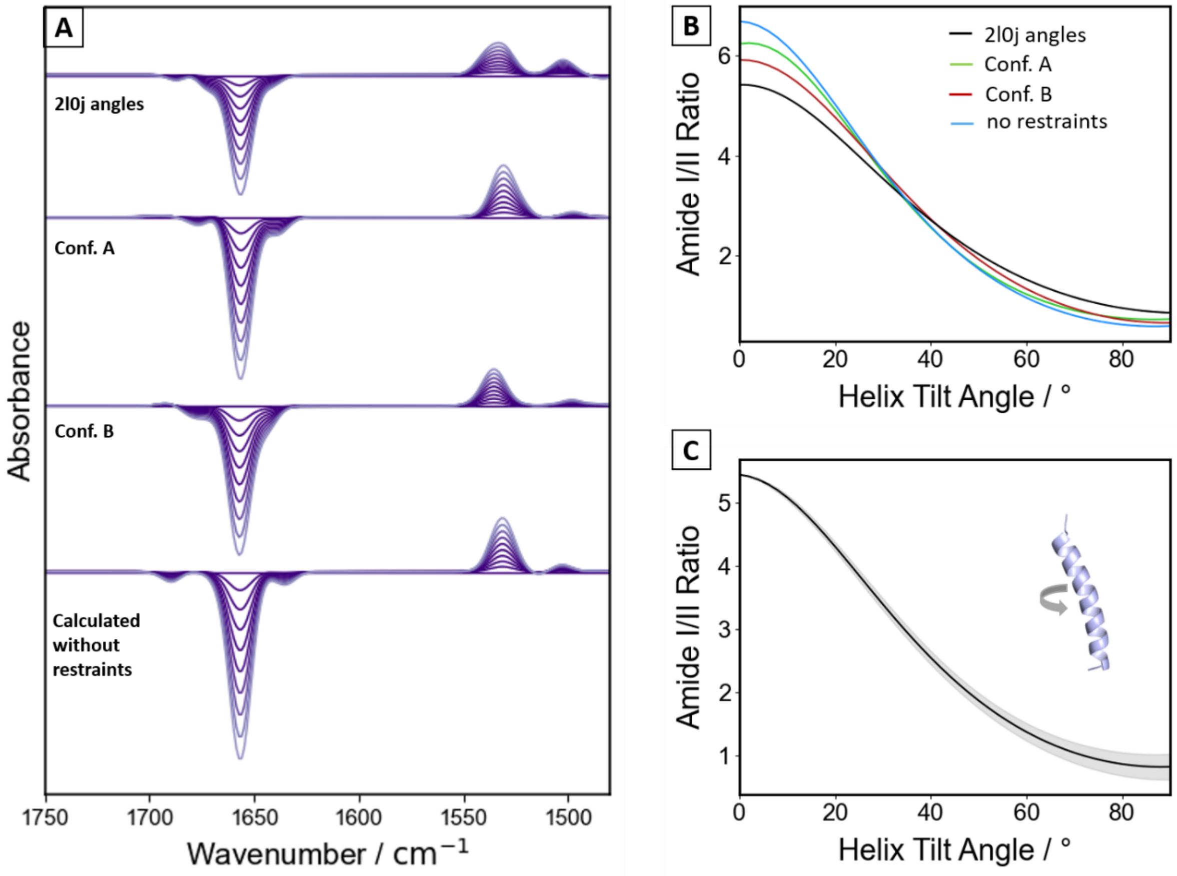

Fig S2: *A: Computational difference spectra for a stepwise TM helix reorientation from 26° to 40°using different dihedral angles as starting structures. B: Trend of the amide I/II ratio of the peak areas dependent on the helix tilt angle in respect to the surface normal from 0° to 90° considering the different starting structures that were calculated and C: considering a difference in the twisting angle of the helix of ±30° from the one that was considered to follow the opening trajectory according to the 6mjh structures. Higher twisting angles led to difference spectra that differed strongly from the experimental results.*

Table S1: *Dihedral angles of the protein backbone derived from different datasets (from 2l0j protein structure and from the TALOS+ analysis of the solid-state NMR data*[8] *(Figure 1B and C, Table S3), where two different conformations were assigned.*

| **Amino acid** | | **2l0j**[14] | | **Conformation A** | | **Conformation B** | |
|---|---|---|---|---|---|---|---|
| **#** | **Code** | **Phi** | **Psi** | **Phi** | **Psi** | **Phi** | **Psi** |
| 25 | P | -54.0 | -47.7 | - | - | -56.3 | -32.4 |
| 26 | L | -64.3 | -47.9 | - | - | -67.5 | -34.0 |
| 27 | V | -62.4 | -42.9 | - | - | -64.3 | -44.6 |
| 28 | V | -60.9 | -51.3 | - | - | -63.9 | -42.1 |
| 29 | A | -61.0 | -42.7 | -62.1 | -35.6 | -58.8 | -47.1 |
| 30 | A | -60.9 | -37.3 | -66.7 | -28.9 | -61.4 | -40.1 |
| 31 | S | -67.1 | -50.9 | -64.0 | -39.1 | -60.8 | -42.6 |
| 32 | I | -49.5 | -47.9 | -64.0 | -38.1 | -60.5 | -45.9 |
| 33 | I | -56.0 | -39.7 | -65.7 | -25.7 | -62.6 | -45.1 |
| 34 | G | -63.8 | -36.4 | -74.7 | -29.2 | - | - |
| 35 | I | -65.7 | -45.8 | -64.4 | -38.9 | - | - |
| 36 | L | -53.7 | -43.5 | -65.6 | -25.2 | - | - |
| 37 | H | -62.2 | -41.9 | -67.0 | -27.6 | - | - |
| 38 | L | -60.0 | -44.6 | -63.1 | -34.9 | - | - |
| 39 | I | -59.9 | -42.2 | -63.6 | -32.3 | - | - |
| 40 | L | -62.5 | -40.9 | -65.7 | -39.3 | - | - |
| 41 | W | -62.9 | -37.6 | -63.2 | -42.9 | - | - |
| 42 | I | -63.3 | -46.1 | -63.2 | -44.6 | - | - |
| 43 | L | -55.0 | -40.6 | -62.0 | -40.5 | - | - |
| 44 | D | -56.6 | -44.4 | -66.7 | -35.0 | - | - |
| 45 | R | -100.7 | -18.4 | -80.0 | -22.4 | - | - |
| 46 | L | -72.7 | -46.6 | -70.8 | -13.5 | - | - |
| 47 | F | -113.5 | 64.5 | -108.4 | 3.4 | - | - |
| 48 | F | -54.2 | -32.4 | 58.0 | 37.3 | - | - |
| 49 | K | -75.4 | -15.8 | -92.5 | -14.6 | - | - |
| 50 | S | -70.6 | -38.7 | -88.2 | -19.8 | - | - |
| 51 | I | -72.8 | -27.9 | -60.7 | -37.4 | - | - |

*Table S2: Tilt angles derived from published M2 structures including experimental parameters.*

| # | PDB ID | Year | author | Seq. | mutation | method | resolution */bb RMSD [Å]* | environment | pH | tilt angle [°] | state |
|---|---|---|---|---|---|---|---|---|---|---|---|
| 1 | 6US8 | 2020 | Thomaston et al. | TM 22-46 | wild type | x-ray diffr. | 1.70 | Lipid Cupic Phase | 7.5 | 23 | S-rimantadine, inward-closed |
| 2 | 6US9 | 2020 | Thomaston et al. | TM 22-46 | wild type | x-ray diffr. | 1.70 | Lipid Cupic Phase | 8.5 | 26 | R-rimantadine, inward-closed |
| 3 | 6NV1 | 2020 | Thomaston et al. | TM 22-46 | V27A | x-ray diffr. | 2.50 | Lipid Cupic Phase | 7.5 | 24 | amine inhibitor, inward-closed |
| 4 | 6OUG | 2020 | Thomaston et al. | 21-61 | V27A | x-ray diffr. | 3.01 | Lipid Cupic Phase | 8.0 | 22 | amine inhibitor, inward-closed |
| 5 | 6MJH | 2019 | Thomaston et al. | TM 22-46 | S31N | x-ray diffr. | 2.06 | lipidic cupic phase, MNG | 5.0 | 17 | closed |
| 5 | 6MJH | 2019 | Thomaston et al. | TM 22-46 | S31N | x-ray diffr. | 2.06 | lipidic cupic phase, MNG | 5.0 | 28 | opened |
| 6 | 6BKK | 2018 | Thomaston et al. | TM 22-46 | wild type | x-ray diffr. | 2.00 | lipidic cubic phase | 5.6 | 26 | amantadine, inward-closed |
| 7 | 6BKL | 2018 | Thomaston et al. | TM 22-46 | wild type | x-ray diffr. | 2.00 | lipidic cubic phase | 4.5 | 26 | rimantadine, inward-closed |
| 8 | 6BOC | 2018 | Thomaston et al. | TM 22-46 | wild type | x-ray diffr. | 2.25 | lipidic cubic phase | 3.5 | 35 | rimantadine, inward-open |
| 9 | 6BMZ | 2018 | Thomaston et al. | TM 22-46 | wild type | x-ray diffr. | 2.63 | lipidic cubic phase | 7.0 | 24 | amine inhibitor, inward-closed |
| 10 | 2LY0 | 2018 | Wang et al. | 19-49 | S31N | sol. NMR | *1.00* | DPC | 6.8 | 22 | M2WJ332 inhibitor |
| 11 | 5JOO | 2017 | Thomaston et al. | TM 22-46 | wild type | XFEL | 1.41 | lipidic cubic phase | 5.5 | 27 | |
| 12 | 5TTC | 2017 | Thomaston et al. | TM 22-46 | wild type | XFEL | 1.40 | lipidic cubic phase | 8.0 | 26 | |
| 13 | 5UM1 | 2017 | Thomaston et al. | TM 22-46 | wild type | XFEL | 1.45 | lipidic cubic phase | 6.5 | 26 | |
| 14 | 5C02 | 2016 | Thomaston, DeGrado | TM 22-46 | S31N | x-ray diffr. | 1.59 | lipidic cubic phase | 8.0 | 25 | |
| 15 | 4QK7 | 2015 | Thomaston et al. | TM 22-46 | wild type | x-ray diffr. | 1.10 | lipidic cubic phase | 8.0 | 29 | |
| 16 | 4QKC | 2015 | Thomaston et al. | TM 22-46 | wild type | x-ray diffr. | 1.10 | lipidic cubic phase | 5.5 | 27 | |
| 17 | 4QKL | 2015 | Thomaston et al. | TM 22-46 | wild type | x-ray diffr. | 1.71 | lipidic cubic phase | 8.0 | 28 | |
| 18 | 4QKM | 2015 | Thomaston et al. | TM 22-46 | wild type | x-ray diffr. | 1.44 | lipidic cubic phase | 5.5 | 25 | |
| 19 | 2N70 | 2015 | Andreas et al. | 18-60 | S31N | ssNMR | *1.10* | lipid bilayer | 7.8 | 20 | |
| 20 | 2MUV | 2014 | Wu et al. | 19-49 | S31N | sol. NMR | *2.50* | DPC micelles | 6.8 | 19 | bound to drug 11 |
| 21 | 2MUW | 2014 | Wu et al. | 19-49 | wild type | sol. NMR | *4.00* | DPC micelles | 7.5 | 19 | bound to drug 11 |
| 22 | 2L0J | 2010 | Sharma et al. | 22-62 | wild type | ssNMR | *0.60* | lipid bilayer DOPE/DOPC | 7.5 | 28 | |
| 23 | 3LBW | 2010 | Acharya et al. | TM 24-46 | G34A | x-ray diffr. | 1.65 | OG | 6.5 | 25 | |
| 24 | 2KQT | 2010 | Cady et al. | TM 22-46 | wild type | ssNMR | *0.67* | DMPC lipid bilayers | 7.5 | 22 | deuterated amantadine |
| 25 | 2KWX | 2010 | Pielak, Chou | 18-60 | V27A | sol. NMR | *0.92* | DHPC | 7.5 | 16 | rimantadine |
| 26 | 2KAD | 2009 | Cady et al. | TM 22-46 | S31N | ssNMR | *n/s* | DLPC bilayer | 7.5 | 38 | with amantadine |
| 27 | 2KIH | 2009 | Pielak et al. | 18-60 | S31N | sol. NMR | *≤1.01* | deuterated DHPC | n/s | 14 | rimantadine |
| 28 | 2RLF | 2008 | Schnell, Chou | 18-60 | wild type | sol. NMR | *≤0.56* | DHPC micelles | 7.5 | 17 | with rimantadine |

| | | | | | | | | | | | |
|---|---|---|---|---|---|---|---|---|---|---|---|
| 29 | 3BKD | 2008 | Stouffer et al. | TM 22-46 | I33Se-Met | x-ray diffr. | 2.05 | BOG, CL, PEG, OG micelles | 7.3 | 30 | |
| 30 | 3C9J | 2008 | Stouffer et al. | TM 22-46 | G34A | x-ray diffr. | 3.50 | OG | 5.3 | 35 | with amantadine |
| 31 | 2H95 | 2007 | Hu et al. | TM 26-43 | wild type | ssNMR | *n/s* | DMPC/DMPG liposomes | 8.8 | 25 | amantadine-blocked |
| 32 | 1NYJ | 2002 | Nishimura et al. | TM 22-46 | wild type | ssNMR | *n/s* | DMPC bilayer | 7.0 | 38 | |

Table S3: Assigned chemical shifts in ppm for M2(18 – 60) from solid-state NMR measurements on the $^{13}C$,$^{15}N$-labeled sample in DPhPC proteoliposomes.

| Residue number | Residue | Atom | Chemical shift conformation A | Chemical shift conformation B |
|---|---|---|---|---|
| 24 | D | C | | |
| 24 | D | CA | | 51.92 |
| 24 | D | CB | | 43.18 |
| 24 | D | N | | 125.56 |
| 25 | P | C | | 177.29 |
| 25 | P | CA | | |
| 25 | P | CB | | |
| 25 | P | N | | |
| 26 | L | C | | 178.32 |
| 26 | L | CA | | 57.15 |
| 26 | L | CB | | 39.74 |
| 26 | L | N | | 118.60 |
| 27 | V | C | | 178.93 |
| 27 | V | CA | | 66.91 |
| 27 | V | CB | | 31.20 |
| 27 | V | N | | 120.44 |
| 28 | V | C | 178.03 | 178.01 |
| 28 | V | CA | | 66.71 |
| 28 | V | CB | | 31.16 |
| 28 | V | N | | 119.66 |
| 29 | A | C | 178.47 | 178.43 |
| 29 | A | CA | 55.36 | 55.56 |
| 29 | A | CB | 18.22 | 18.32 |
| 29 | A | N | 119.93 | 121.21 |
| 30 | A | C | 178.47 | 178.33 |
| 30 | A | CA | 54.89 | 54.66 |
| 30 | A | CB | 18.06 | 18.20 |
| 30 | A | N | 117.63 | 118.16 |
| 31 | S | C | 175.20 | 175.38 |
| 31 | S | CA | 62.31 | 62.52 |
| 31 | S | CB | 65.38 | 65.54 |
| 31 | S | N | 112.54 | 112.89 |
| 32 | I | C | 177.05 | 177.42 |
| 32 | I | CA | 63.25 | 65.40 |
| 32 | I | CB | 37.03 | 37.56 |
| 32 | I | N | 120.26 | 119.99 |
| 33 | I | C | 176.99 | 177.35 |
| 33 | I | CA | 65.30 | 65.87 |
| 33 | I | CB | 37.00 | 36.73 |
| 33 | I | N | 118.76 | 114.51 |
| 34 | G | C | 175.11 | |
| 34 | G | CA | 48.07 | 47.52 |

| | | | | |
|---|---|---|---|---|
| 34 | G | N | 105.33 | 106.63 |
| 35 | I | C | 176.85 | |
| 35 | I | CA | 65.14 | |
| 35 | I | CB | 37.24 | |
| 35 | I | N | 119.79 | |
| 36 | L | C | 177.66 | |
| 36 | L | CA | 57.99 | |
| 36 | L | CB | 41.46 | |
| 36 | L | N | 119.43 | |
| 37 | H | C | 175.16 | |
| 37 | H | CA | 59.12 | |
| 37 | H | CB | 30.25 | |
| 37 | H | N | 116.59 | |
| 38 | L | C | 177.52 | |
| 38 | L | CA | 58.61 | |
| 38 | L | CB | 42.05 | |
| 38 | L | N | 117.40 | |
| 39 | I | C | 177.14 | |
| 39 | I | CA | 64.86 | |
| 39 | I | CB | 37.44 | |
| 39 | I | N | 114.28 | |
| 40 | L | C | 178.69 | |
| 40 | L | CA | 58.28 | |
| 40 | L | CB | 40.75 | |
| 40 | L | N | 121.62 | |
| 41 | W | C | 178.27 | |
| 41 | W | CA | 61.11 | |
| 41 | W | CB | 27.50 | |
| 41 | W | N | 120.76 | |
| 42 | I | C | 177.85 | |
| 42 | I | CA | 66.23 | |
| 42 | I | CB | 37.12 | |
| 42 | I | N | 117.39 | |
| 43 | L | C | 179.68 | |
| 43 | L | CA | 57.70 | |
| 43 | L | CB | 41.49 | |
| 43 | L | N | 117.27 | |
| 44 | D | C | 178.35 | |
| 44 | D | CA | 57.57 | |
| 44 | D | CB | 42.13 | |
| 44 | D | N | 120.56 | |
| 45 | R | C | 178.50 | |
| 45 | R | CA | 56.67 | |
| 45 | R | CB | 29.76 | |
| 45 | R | N | 116.56 | |
| 46 | L | C | 177.45 | |

| | | | |
|---|---|---|---|
| 46 | L | CA | 56.03 |
| 46 | L | CB | 41.77 |
| 46 | L | N | 113.39 |
| 47 | F | C | 175.61 |
| 47 | F | CA | 58.48 |
| 47 | F | CB | |
| 47 | F | N | 112.53 |
| 48 | F | C | 175.93 |
| 48 | F | CA | 62.24 |
| 48 | F | CB | 31.26 |
| 48 | F | N | 117.26 |
| 49 | K | C | 175.59 |
| 49 | K | CA | 56.83 |
| 49 | K | CB | 30.04 |
| 49 | K | N | 120.46 |
| 50 | S | C | 174.76 |
| 50 | S | CA | 60.10 |
| 50 | S | CB | 66.08 |
| 50 | S | N | 111.19 |
| 51 | I | C | 176.44 |
| 51 | I | CA | 65.60 |
| 51 | I | CB | 37.89 |
| 51 | I | N | 126.67 |
| 52 | Y | C | 178.94 |
| 52 | Y | CA | 62.61 |
| 52 | Y | CB | |
| 52 | Y | N | 117.31 |
